# Sex differences in brain-behavior relationships in the first two years of life

**DOI:** 10.1101/2024.01.31.578147

**Authors:** Sonja J. Fenske, Janelle Liu, Haitao Chen, Marcio A. Diniz, Rebecca L. Stephens, Emil Cornea, John H. Gilmore, Wei Gao

**Author notes:** Corresponding Author: Wei Gao, Ph.D., Biomedical Imaging Research Institute, Department of Biomedical Sciences and Imaging, Cedars–Sinai Medical Center, 116 N. Robertson Blvd., PACT 800.7G, Los Angeles, CA 90048.

## Abstract

**Background:** Evidence for sex differences in cognition in childhood is established, but less is known about the underlying neural mechanisms for these differences. Recent findings suggest the existence of brain-behavior relationship heterogeneities during infancy; however, it remains unclear whether sex underlies these heterogeneities during this critical period when sex-related behavioral differences arise.

**Methods:** A sample of 316 infants was included with resting-state functional magnetic resonance imaging scans at neonate (3 weeks), 1, and 2 years of age. We used multiple linear regression to test interactions between sex and resting-state functional connectivity on behavioral scores of working memory, inhibitory self-control, intelligence, and anxiety collected at 4 years of age.

**Results:** We found six age-specific, intra-hemispheric connections showing significant and robust sex differences in functional connectivity-behavior relationships. All connections are either with the prefrontal cortex or the temporal pole, which has direct anatomical pathways to the prefrontal cortex. Sex differences in functional connectivity only emerge when associated with behavior, and not in functional connectivity alone. Furthermore, at neonate and 2 years of age, these age-specific connections displayed greater connectivity in males and lower connectivity in females in association with better behavioral scores.

**Conclusions:** Taken together, we critically capture robust and conserved brain mechanisms that are distinct to sex and are defined by their relationship to behavioral outcomes. Our results establish brain-behavior mechanisms as an important feature in the search for sex differences during development.

**Plain language summary:** Early childhood differences exist in mental processes and behavior between males and females. The brain-basis for these sex differences may arise in infancy. Indeed, small brain differences in infancy may contribute to major changes in cognitive ability throughout childhood. However, few studies have examined sex differences in brain functionality in infancy and their relationship to future behaviors in early childhood. In this study, we aimed to study this relationship by using sex differences in brain functional measures in neonate, 1, and 2-year-olds and 4-year behavioral outcomes. We identified six functional connections with robust brain-behavior sex differences. These connections were unique to frontal brain regions. Also, these connections were not specific to the brain and were only evident when associated with future behavior. In brief, our analysis shows distinct age-specific brain-behavior relationships in males and females in early childhood. This is helpful for a better understanding of brain-based prediction of behavior and informed intervention of future disorders and disabilities characterized by a sex bias.

**Highlights:**

- Multiple linear regression was used to test the interaction between sex and early childhood resting-state functional connectivity on future behavioral scores
- Six age-specific, intra-hemispheric functional connections displayed sex differences
- Most connections exist within prefrontal regions (with one connection in the temporal pole)
- Functional connections are specific to brain-behavior relationships and not in brain connectivity alone
- Sex differences in brain-behavior relationships are robust at smaller sample sizes

## 1. Introduction

Sex differences in cognition and behavior during early development are well documented. For example, females tend to perform better than males in measures of executive functioning [1,2], and intelligence [3,4] in toddler and preschool years. Previous studies suggest that sex differences in these cognitive processes may arise from changes in developmental trajectories [5,6] with females maturing earlier. Even so, these processes correspond with sex differences seen throughout childhood and adolescence in reading comprehension [5,7], language production [5,8,9], working memory [10–12], and socioemotional processing [11,13,14]. Clinically, deficits in executive functioning and intelligence can serve as markers for the diagnoses of certain neurodevelopmental disorders and learning disabilities, such as autism (ASD) and attention deficit hyperactivity disorder (ADHD). Diagnoses of ASD and ADHD also have a higher prevalence in males [15–19] which may occur as a result of predominantly male-derived criteria [20–24]. These discrepancies in sex may lead to a misdiagnosis in females, delaying or preventing treatment [20–22]. However, the neural underpinnings of these sex differences in cognition remain poorly delineated.

Previous studies have focused on sex differences in the brain using resting-state functional magnetic resonance imaging (rsfMRI). In older children and adults, sex differences in brain structure and function are sparse and small [25–28]. An even smaller number of infant studies have investigated and shown sex differences in rsfMRI [28]. Our past work identified significant sex differences in the rate of growth of the frontoparietal network from birth to 2 years of age [29], as well as female-driven global changes in connectivity in the first year of life [30]. Our most recent study found consistent sex differences in the temporal areas of the brain in the first two years of life [31]. Although we examined relationships between temporal functional connectivity and behavioral scores, whole-brain brain-behavior relationships were not a focus this of previous work. By investigating sex differences in the brain exclusively, previous studies may have missed an important source of variation only seen in mechanisms underlying brain-behavior relationships.

Even if similarities in brain structure and functional organization are observed during development, subpopulations can develop highly divergent behavioral outputs. This divergence may be a result of different brain-behavioral mechanisms employed by each subpopulation. For example, despite many sex differences in the brain showing a small or negligible effect [25,26,32–34], such differences are not necessarily trivial and may, in fact, have a meaningful behavioral consequence [33–35]. Indeed, small brain differences in infancy may contribute to major changes in cognitive ability throughout childhood and adulthood [28,36–39]. However, studies demonstrating these longitudinal brain-behavior relationships are limited by small effect size. We previously identified subgroups of infants with similar neonatal functional connectivity but divergent future behavioral scores of intelligence as well as contrasting brain-behavior relationships [36]. However, the role of sex in this data-driven subgrouping of infants remains to be studied. In adults, Ragland et al. reported no main effect of sex in their whole-brain analysis but found brain-behavior relationships between resting regional cerebral blood flow and verbal episodic memory after a post hoc analysis examining correlations in males and females separately [40]. Moreover, sex differences in the brain can be lost at group-level statistics which fails to capture intersubject variability [41,42]. Ultimately, suitable identification of underlying neural mechanisms rests on the principle that we consider sex-based heterogeneity in brain-behavior relationships [36].

To this end, we investigated whether the relationship between whole-brain functional connectivity measured in infancy and early childhood behavioral responses differed between males and females. The misreporting of significant within-group results (i.e., testing males and females separately) as statistically different between-group results (i.e., males vs. females) [43–45] has been a critique in the analysis of sex differences in the biological sciences. Instead, the use of multiple linear regression to test interaction effects between sex and the variable of interest is recommended. In this study, we tested between-group differences in infant rsfMRI functional connectivity measures and behavioral scores of working memory, inhibitory self-control, intelligence, and anxiety collected at 4 years of age. We hypothesized that functional connections in infancy associated with future behavioral outcomes in childhood would differ due to sex.

## 2. Methods

### 2.1 Infant participants and imaging acquisition

Data were collected from typically developing infants as part of the University of North Carolina Early Brain Development Study to investigate early childhood brain and behavior [28,46]. Parents or legal guardians of infants provided informed consent under protocols sanctioned by the University of North Carolina at Chapel Hill and Cedars-Sinai Institutional Review Boards. After quality control, we retrospectively identified 316 participants with at least one successful rsfMRI scan in the first two years of life. We included timepoints at neonate (3 weeks from birth) (*n*=229), 1 year (*n*=146), and 2 years (*n*=109) of age. Twin status was included as a control variable. We treated gestational age at birth <37 weeks and maternal psychiatric disorder status as exclusionary criteria. Participant cohort characteristics are defined in Table 1.

**Table 1:**
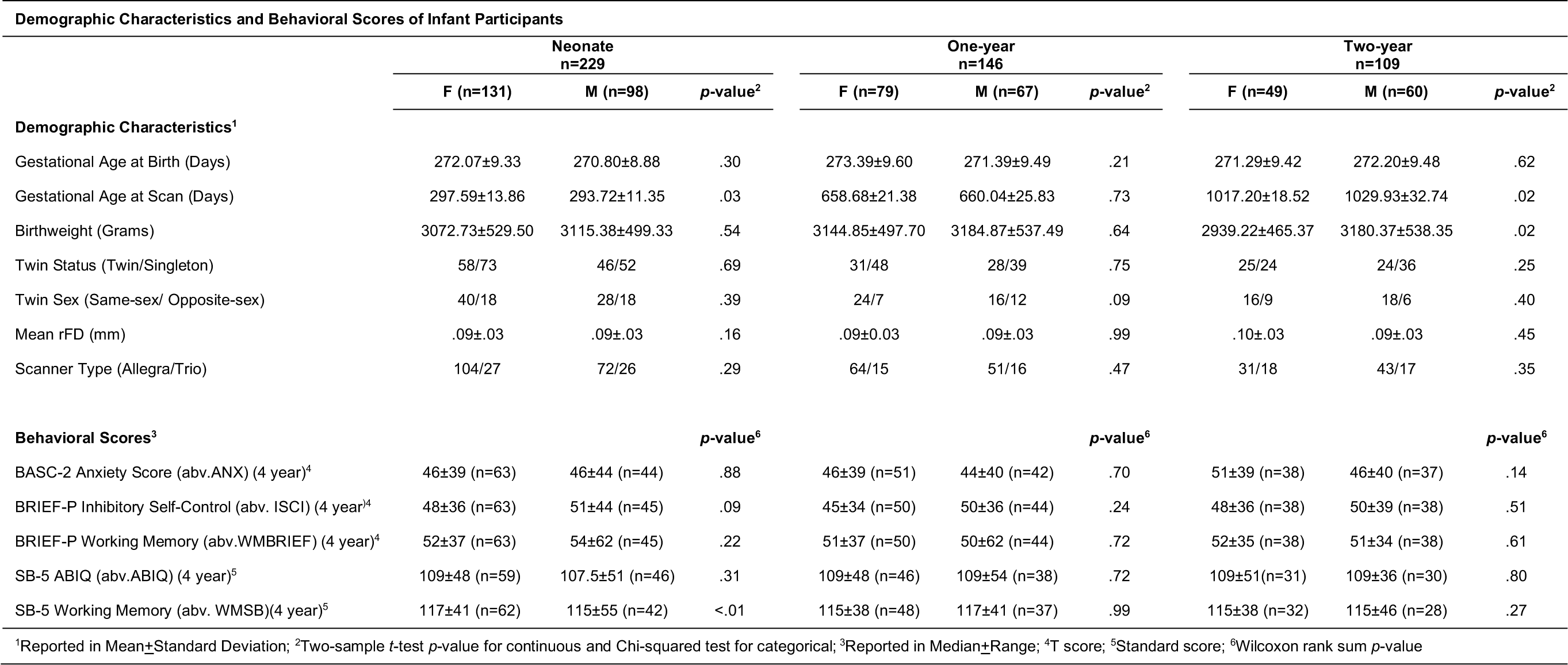
Participant demographic characteristics and 4-year behavioral scores. Key: Female (F), Male (M), Behavior Assessment System for Children – Second Edition (BASC-2), Behavior Rating Inventory of Executive Function – Preschool version (BRIEF-P), and Stanford-Binet Intelligence Scales – Fifth Edition (SB-5).

The magnetic resonance imaging (MRI) collection follows the same procedure and data acquisition procedure for previous studies [30,31,36]. Participants were scanned during natural sleep. The distribution and retention of longitudinal data collected at neonates, 1 year, and 2 years of age are displayed in Supplementary Figure 1. Attrition across timepoints did not influence differences in sex (*p*=.12) or race (*p*=.08). MRI data were collected on a Siemens 3T Allegra (neonate: n=176; 1-year-old: *n*=115; 2-year-old: *n*=74) or TIM Trio (neonate: *n*=53; 1-year-old: *n*=31; 2-year-old: *n*=35) scanner with a circularly polarized head coil or a 32-channel head coil, respectively. We included scanner as a control variable in our subsequent analysis. Functional images were acquired using a T2*-weighted echo planner imaging (EPI) sequence: TR=2000ms, TE=32ms, 33 slices, 150 volumes, 4mm^3^ voxel size, and anterior-posterior phase encoding direction. Structural images were acquired using a three-dimensional magnetization prepared rapid acquisition gradient-echo (MPRAGE) sequence: TR=2000ms, TE=4.38ms, inversion time=1100ms, and 1mm^3^ voxel size.

### 2.2 Image preprocessing

Our rsfMRI preprocessing pipeline was developed in prior work [31,36,38,47,48] using FMRIB’s Software Library (FSL, version 6.0, [49,50]), Advanced Normalization Tools (ANTs, version 1.10.11, [51]), and Analysis of Functional Neuroimaging (AFNI, Version AFNI_19.3.12, [52]). Preprocessing steps included ridged-body motion correction, scrubbing, co-registration, nuisance signal regression, bandpass filtering (0.01-0.08 Hz), registration, spatial smoothing, and global signal regression. Motion-related nuisance variables included 24 parameters (six motion estimates and their derivative, quadratic, and squared derivative terms [53]). Mean white matter and mean cerebrospinal fluid as well as their temporal derivatives and quadratic terms were also included as nuisance variables. Bandpass filtering was performed after regression of the nuisance variables on the functional dataset. Data scrubbing was applied by removing volumes with frame-wise displacements (FD) > 0.3mm. FD was calculated following motion correction and these volumes were interpolated in the functional time series. After bandpass filtering these volumes were removed (“scrubbed”). The scrubbed functional datasets were non-linearly registered to age-specific and 2-year templates at 4 mm isotropic resolution [54] (ANTs rigid + affine + deformable registration, version 1.10.11, [51]). The 2-year template served as the final target for spatial registration for all timepoints (i.e., neonates, 1-, and 2-year-olds). Spatial smoothing was performed using a Gaussian kernel of 6 mm full width at half maximum (FWHM). The functional image was then truncated to 90 volumes. Participants with <90 volumes remaining (3 minutes) were excluded (neonate: *n*=8; 1-year-old: *n*=1; 2-year-old: *n*=0). Finally, each participant’s mean global time-series signal from the grey matter was regressed from the functional data.

### 2.3 Behavioral scores

In this study, we obtained five measures of parent-report and task-based laboratory assessments of behavior collected at 4 years of age. These 4-year behavioral scores have been previously described in detail [31,38,55]. Measures of anxiety were assessed using the Behavior Assessment System for Children – Second Edition (BASC-2; [56]). This parent-report assessment was designed to test behaviors such as “worry or fear about real or imagined problems” [38]. Indices of working memory (WM) and inhibitory self-control (ISCI) were assessed using the Behavior Rating Inventory of Executive Function – Preschool version (BRIEF-P, [57]). The parent-report assessment for WM were designed to test “the capacity to hold information… [to] complete a task, encode information, or generating goals, plans, and sequential steps to achieving goals” [57]. The parent-report assessment for ISCI measured “the ability to regulate behavioral and emotional responses across different situations” [38]. We also used subscales of intelligence (ABIQ) and WM from the Stanford-Binet Intelligence Scales – Fifth Edition (SB-5, [58]). These task-based assessments for ABIQ tested general cognitive ability, and assessments for WM examined “the ability to store, transform and retrieve information from short-term memory stores” [55,59]. All measures have robust test-retest reliability [36,38,57,59,60]. Higher scores on BASC-2 and BRIEF-P assessments indicate more problematic behavioral outcomes, whereas higher scores on the SB-5 indicate better behavioral outcomes.

### 2.4 Experimental Design and statistical analysis

#### 2.4.1 Whole-brain connectivity

We quantified functional measures throughout the brain at each infant timepoint. The residual blood oxygen level-dependent (BOLD) time-series was extracted and averaged using the 278 functional parcellations from the 2-year UNC Cedars Infant Atlas [61]. For each participant, a 278×278 functional connectivity matrix was created using the pair-wise correlation between the average signal from each region of interest (ROI) and every other ROI across the whole-brain. Correlation measures were normalized using Fisher-Z transformation. The upper triangular, discarding the diagonal, was taken from each functional connectivity matrix. These steps provided a total of 38,503 unique connections per participant.

#### 2.4.2 Detection of brain-behavior relationships

We used multiple linear regression to test significant interactions between sex and whole-brain functional connectivity on future behavioral scores taken at 4 years of age. To maintain consistency with previously published work [36,47,48], we controlled for the following participant characteristics: gestational age at birth, gestational age at scan (scaled by the natural logarithm), birthweight, twin status, mean rFD, and scanner. Maternal education did not affect the significance of interactions and so was excluded as a control variable. We adjusted for these control variables in our linear regression at each functional connection. To detect significant interactions, a stringent threshold of *p*≤1.299×10^-6^ was calculated by utilizing Bonferroni correction for multiple comparisons across all connections (*N*=38,503). Throughout this manuscript, the term ‘interaction’ is in reference to the value of the interaction term (sex*functional connectivity) in the linear regression.

To determine if sex differences are specific to brain-behavior relationships, we tested between-group connectivity differences from the connections found in the whole-brain analysis in a post-hoc analysis. First, functional connectivity was calculated within-group across all participants for each timepoint using a Wilcoxon Signed Rank Test (MATLAB’s signrank, *p*≤.05). For example, if an interaction between sex and functional connectivity on behavior was found in 1-year-olds, the functional connectivity from the same ROIs was then tested cross sectionally all neonate, 1-year-old, and 2-year-old males and females, independent of behavior. Second, we tested between-group differences in functional connectivity in these same ROIs at each timepoint using a two-sided Wilcoxon rank sum test (MATLAB’s rank sum, *p*≤.05).

Multiple linear regression was applied to stratified male and female behavior scores to decompose significant interactions. We adjusted for the same control variables in our linear regression model. The resulting residuals were used in Spearman’s rank-order correlation (*p*≤.05) to male and female participants’ functional connectivity. These values were calculated to evaluate the linear relationship between connectivity and behavior in males and females independently.

#### 2.4.3 Consistency testing

We measured the consistency of sex differences by examining brain-behavior relationships across a range of subsamples of the dataset. Specifically, we resampled without replacement (*N*=1000), at five different subsamples (i.e., 90, 80, 70, 60, 50, 40, and 30 percent of the total number of participants). At each subsample, the same multiple linear regression model used to test interactions in our whole-brain analysis was applied to the connections found to have significant brain-behavior sex differences after passing Bonferroni correction. Consistency was defined as the probability of replicating an interaction at a threshold of *p*≤.01. This procedure assessed the stability of the interaction between sex and functional connectivity on 4-year behavioral scores in discrete ROIs at each timepoint.

## 3. Results

### 3.1 Highly significant sex differences in brain-behavior relationships primarily reside within prefrontal regions

Across 38,503 whole-brain functional connections, six age-specific interactions passed Bonferroni corrections for multiple comparisons. Figure 1 shows distributions of uncorrected p-values for the interaction effect across all unique connections associated with a behavioral score for each timepoint (i.e., neonates, 1-, and 2-year-olds). Figure 2 presents a visualization of the significant functional connections. Most connections are located in the prefrontal cortex. Another connection is in the temporal pole, a region that is strongly interconnected with the orbitofrontal cortex [62].

**Figure 1:**
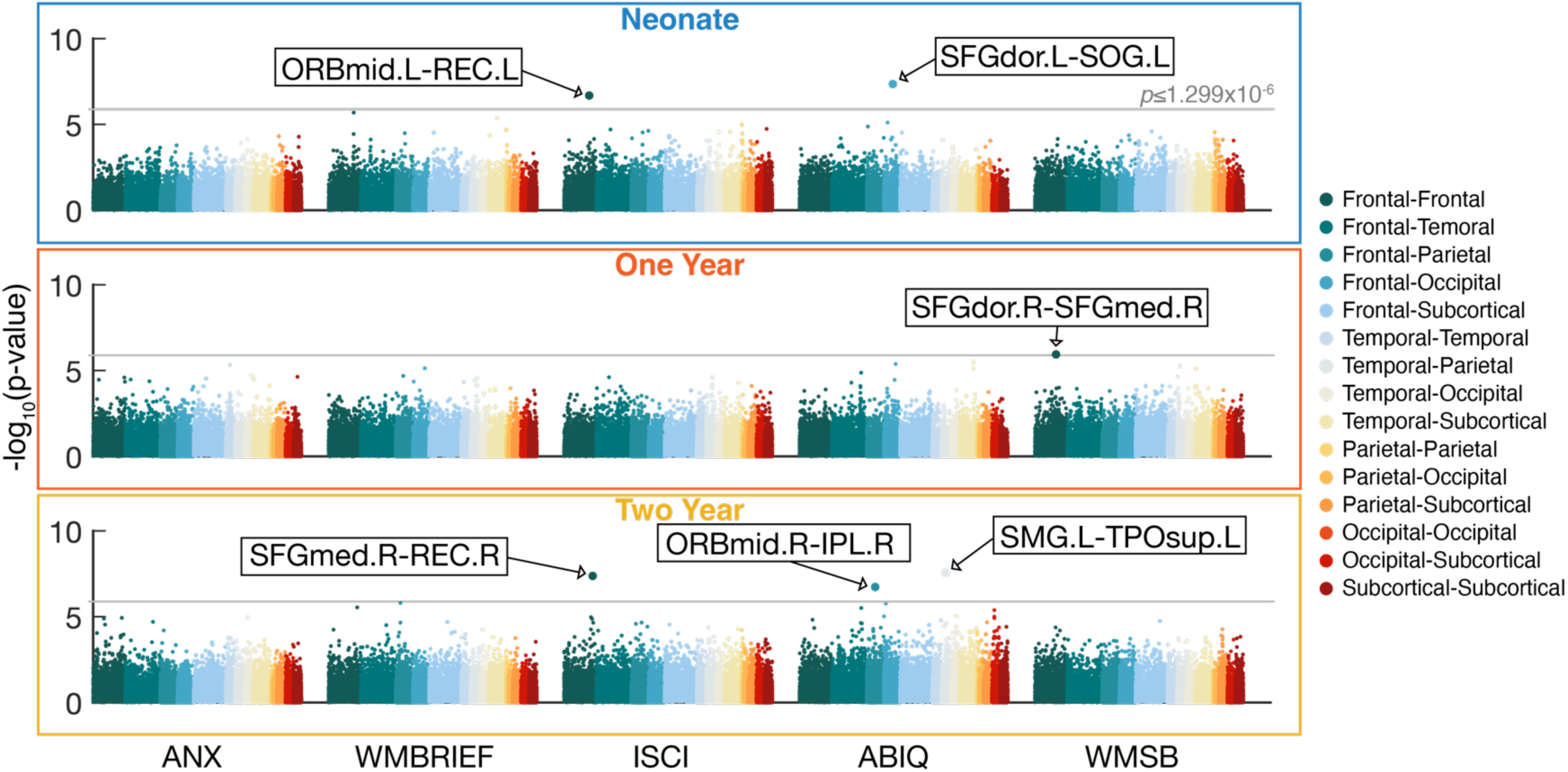
Significant interactions between sex and functional connectivity on 4-year behavioral scores after Bonferroni correction. Scatter plots display the distribution of uncorrected p*-*values for sex and functional connectivity interactions associated with 4-year behavioral scores in neonates, 1-, and 2-year-olds. The value on the y-axis represents the −log_10_ of the p-values. The grey line represents the Bonferroni threshold (*p*≤1.299×10^-6^, -log10(*p*) = 5.89). The resulting six connections are defined by their automated anatomical atlas labels (AAL) from the UNC Cedars Infant Atlas [61]. Key: *IPL*, Inferior parietal lobule, *ORBmid,* Orbitofrontal cortex (middle), *REC*, Rectus gyrus, *SFGdor*, Superior frontal gyrus (dorsal), *SFGmed*, Superior frontal gyrus (medial), *SMG,* Supramarginal gyrus, *SOG*, Superior occipital gyrus, *TPOsup*, Temporal pole (superior); Behavioral scores: *ANX*, BASC-2 Anxiety Score, *WMBREIF*, BRIEF-P Working Memory, *ISCI*, BRIEF-P Inhibitory Self-Control, *ABIQ*, SB-5 Abbreviated Battery IQ, *WMSB*, SB-5 Working Memory.

**Figure 2:**
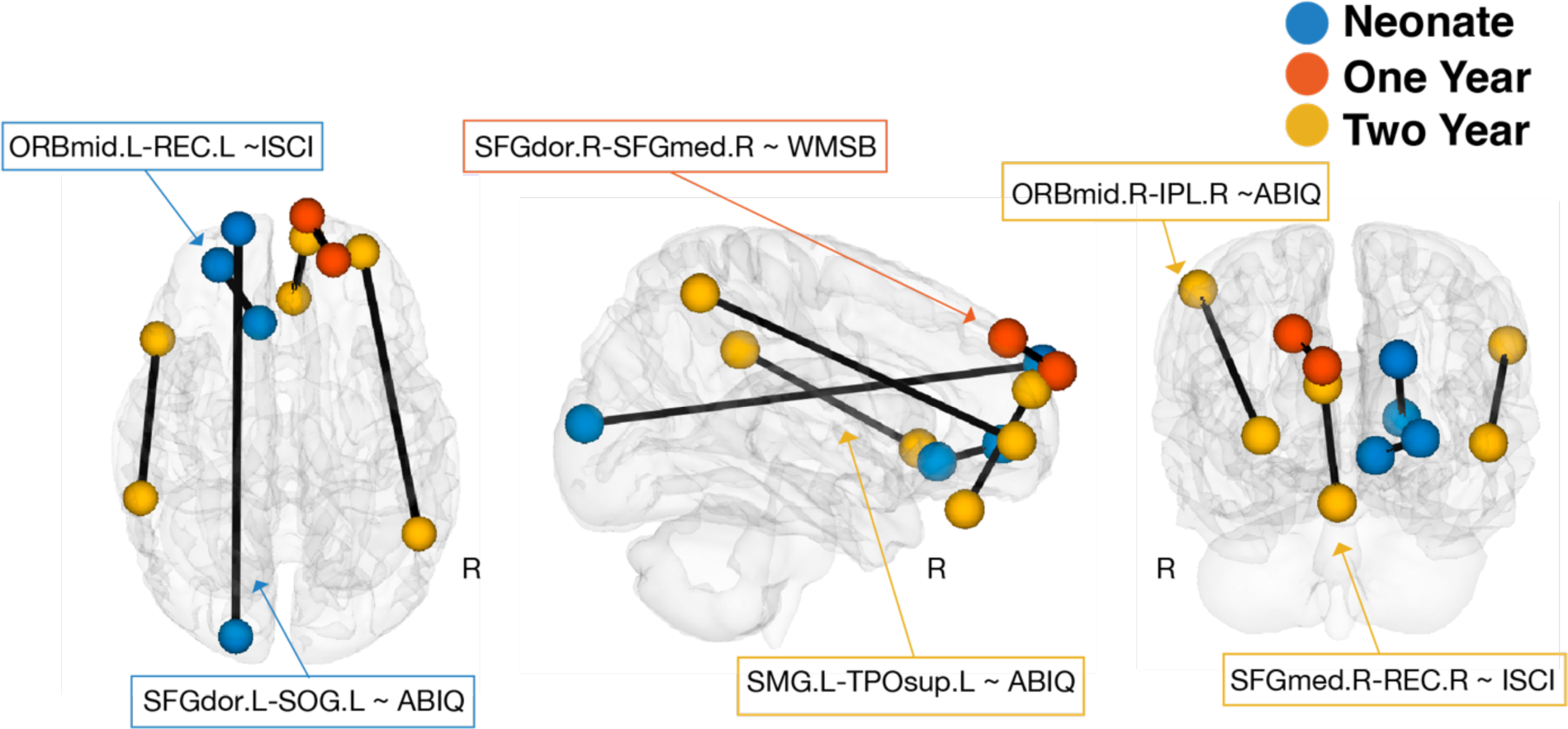
Sex differences in functional connections associated with behavior are mainly located between prefrontal regions to other regions throughout the infant brain. Colored centroids represent ROIs for each of the six connections (defined by their sex*functional connectivity interactions with 4-year behavioral scores, see Figure 1). Key: Neonate: Blue; One Year: Orange; Two Year: Yellow.

All functional connections showing significant interactions were intra-hemispheric. Specifically, we found significant interactions between sex and the left mid-orbitofrontal cortex and the left rectus gyrus in neonates in association with inhibitory self-control collected at 4 years of age (ISCI, *B*=38.43, *df*=98, *p*=2.07×10^-7^), and the left dorsal superior frontal gyrus and the left superior occipital gyrus in neonates in association with intelligence collected at 4 years of age (ABIQ, *B*=-39.21, *df*=95, *p*=4.44×10^-8^); the right dorsal superior frontal gyrus and the right medial superior frontal gyrus in 1-year-olds in association with working memory collected at 4 years of age (WMSB, *B*=35.60, *df*=75, *p*=1.14×10^-6^); the right medial superior frontal gyrus and the right rectus gyrus in 2-year-olds in association with inhibitory self-control collected at 4 years of age (ISCI, *B*=41.20, *df*=66, *p*=4.21×10^-8^), and the left supramarginal gyrus and the left superior temporal pole in 2-year-olds in association with intelligence collected at 4 years of age (ABIQ, *B*=-49.99, *df*=51, *p*=2.79×10^-8^) and the right mid-orbitofrontal cortex and the right inferior parietal lobule in 2-year-olds in association with intelligence collected at 4 years of age (ABIQ, *B*=-50.41, *df*=51, *p*=1.85×10^-7^) in 2-year-olds. These interactions did not pass Bonferroni multiple-comparisons corrections at other timepoints.

### 3.2 Connections showing significant sex differences in brain-behavior relationships failed to show any significant differences when examined on functional connectivity alone

The average within-group connectivity in the six connections showed significance in males and females separately across timepoints (Fig. 3A). The connections that were not significant included: the right orbitofrontal cortex and the right inferior parietal lobule in neonate males; the right medial superior frontal gyrus and the right rectus gyrus in 1-year-old and 2-year-old females; and the left dorsal superior frontal gyrus and the left superior occipital gyrus in 2-year-old males and females.

**Figure 3:**
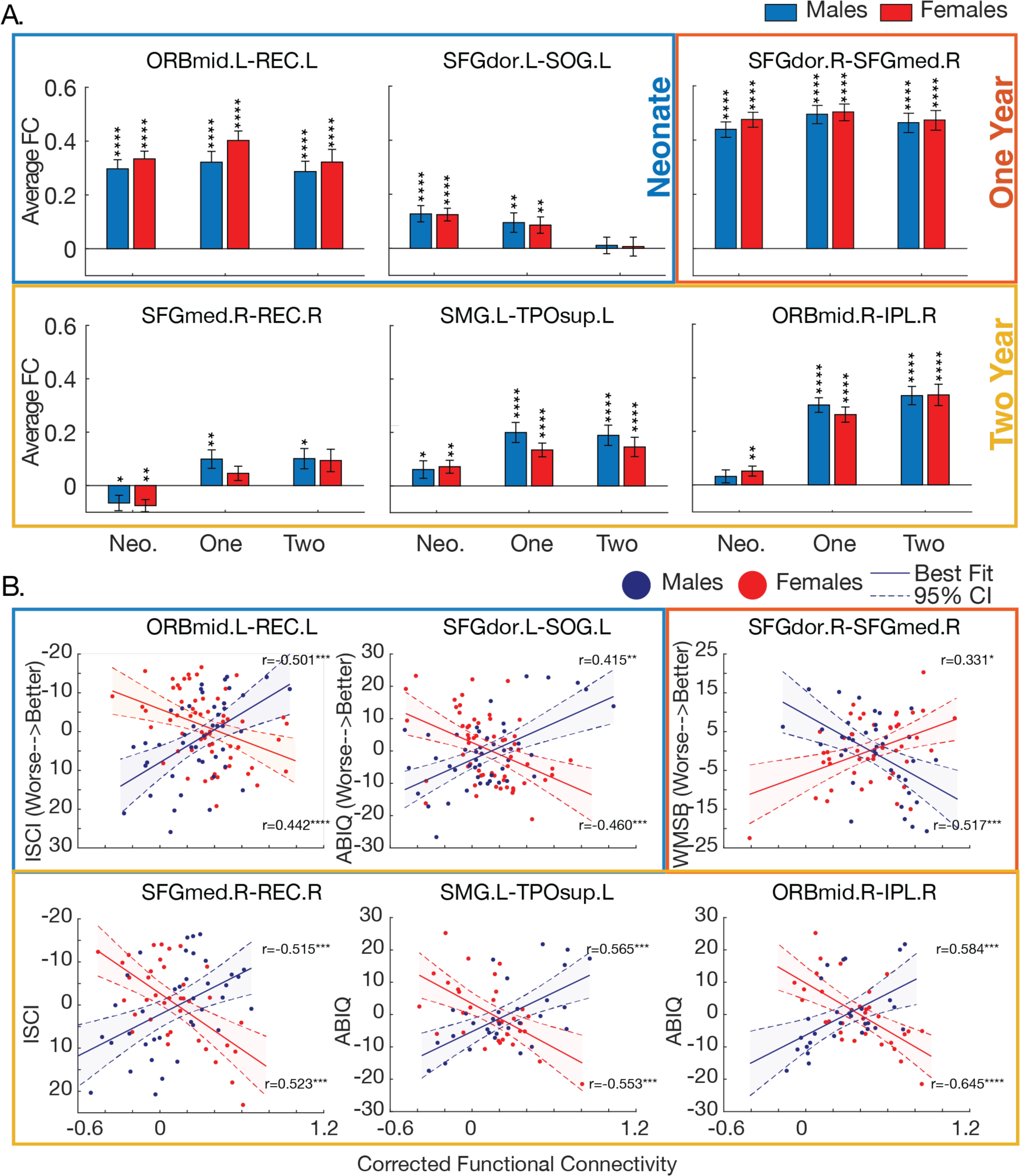
Sex differences in functional connectivity are apparent only when linearly related to behavior. A. Sex differences are not significant in between-group functional connectivity across infancy. Average connectivity of males and females from the six connections across infant ages show significance (*p*≤.05) within males and females independently, but not between males and females. B. Male and female functional connections are linearly related to 4-year behavioral scores. Lines indicate the best linear fit and 95% confidence interval (CI) for males and females separately. Rho values show the strength of the association between connectivity and behavior in either males or females. Asterisks show the significance of the association between connectivity and behavior in either males or females. Better behavioral scores are associated with lower connectivity in females and greater connectivity in males (excl. the brain-behavior relationship in 1-year-olds with opposite effects). The y-axis for ISCI is flipped such that higher scores indicate better outcomes. The colored boxes show each age from which the functional connection was derived. Key: Neonate: Blue; One Year: Orange; Two Year: Yellow.

However, none of the six connections displayed between-group sex differences across timepoints (Fig. 3A). Sex differences in functional connectivity emerge only when these measures are associated with 4-year behavioral scores.

We further examined within-group brain-behavior relationships (Fig. 3B). In neonates, interactions between sex and connectivity showed a similar relationship: greater connectivity in males (M) and lower connectivity in females (F) within 1) the left mid-orbitofrontal cortex and the left rectus gyrus, and 2) the left dorsal superior frontal gyrus and the left superior occipital gyrus were associated with better scores in inhibitory self-control (M: *r*=-0.50 *p*≤.001, F: *r*=0.44 *p*≤.001) and in intelligence (M: *r*=0.42 *p*=.004, F: *r*=-0.46 *p*≤.001), respectively. In 1-year-olds, lower connectivity in males and greater connectivity in females within the right dorsal superior frontal gyrus and the right medial superior frontal gyrus were associated with better scores in working memory (M: r=-0.52 *p*≤.001, F: *r*=0.33 *p*=.02). In 2-year-olds, greater connectivity in males and lower connectivity in females within 1) the right medial superior frontal gyrus and the right rectus gyrus, 2) the left supramarginal gyrus and the left superior temporal pole, and 3) the right mid-orbitofrontal cortex and the right inferior parietal lobule were associated with better behavioral scores in inhibitory self-control (M: *r*=-0.52 *p*≤.001, F: *r*=0.523 *p*≤.001) and intelligence (M: *r*=0.57 *p*≤.001; F: =-0.55, *p*≤.001; M: *r*=0.58, *p*≤.001, F: *r*=-0.65 *p*≤.001), respectively. Apart from 1-year-old connectivity, these age-specific, intra-hemispheric connections displayed greater connectivity in males and lower connectivity in females in association with better behavioral scores.

### 3.3 Significant sex differences in brain-behavior relationships are highly robust against smaller sample sizes

Sampling without replacement (*N*=1000) showed consistent brain-behavior sex differences across smaller subsamples of the original dataset (Fig. 4). In subsamples reduced to 40% of participants, the probability of replicating an interaction effect at a threshold of *p*≤.01 was above chance across all connections. Here, the number of participants in each brain-behavior relationship include: in neonates, *n*=43(25F) for the left mid-orbitofrontal cortex and the left rectus gyrus associated with inhibitory self-control, and *n*=42(25F) for the left dorsal superior frontal gyrus and the left superior occipital gyrus associated with intelligence; in 1-year-olds, *n*=34(20F) for the right dorsal superior frontal gyrus and the right medial superior frontal gyrus associated with working memory; and in 2-year-olds, *n*=23(15F) for the right medial superior frontal gyrus and the right rectus gyrus associated with inhibitory self-control, and *n*=18(12F) for the left supramarginal gyrus and the left superior temporal pole associated with intelligence, and the right mid-orbitofrontal cortex and the right inferior parietal lobule associated with intelligence. To demonstrate the strength of the interaction at each of the six connections, we randomly pulled one iteration of subsampled data at 40% of participants (see Fig. 4B).

**Figure 4:**
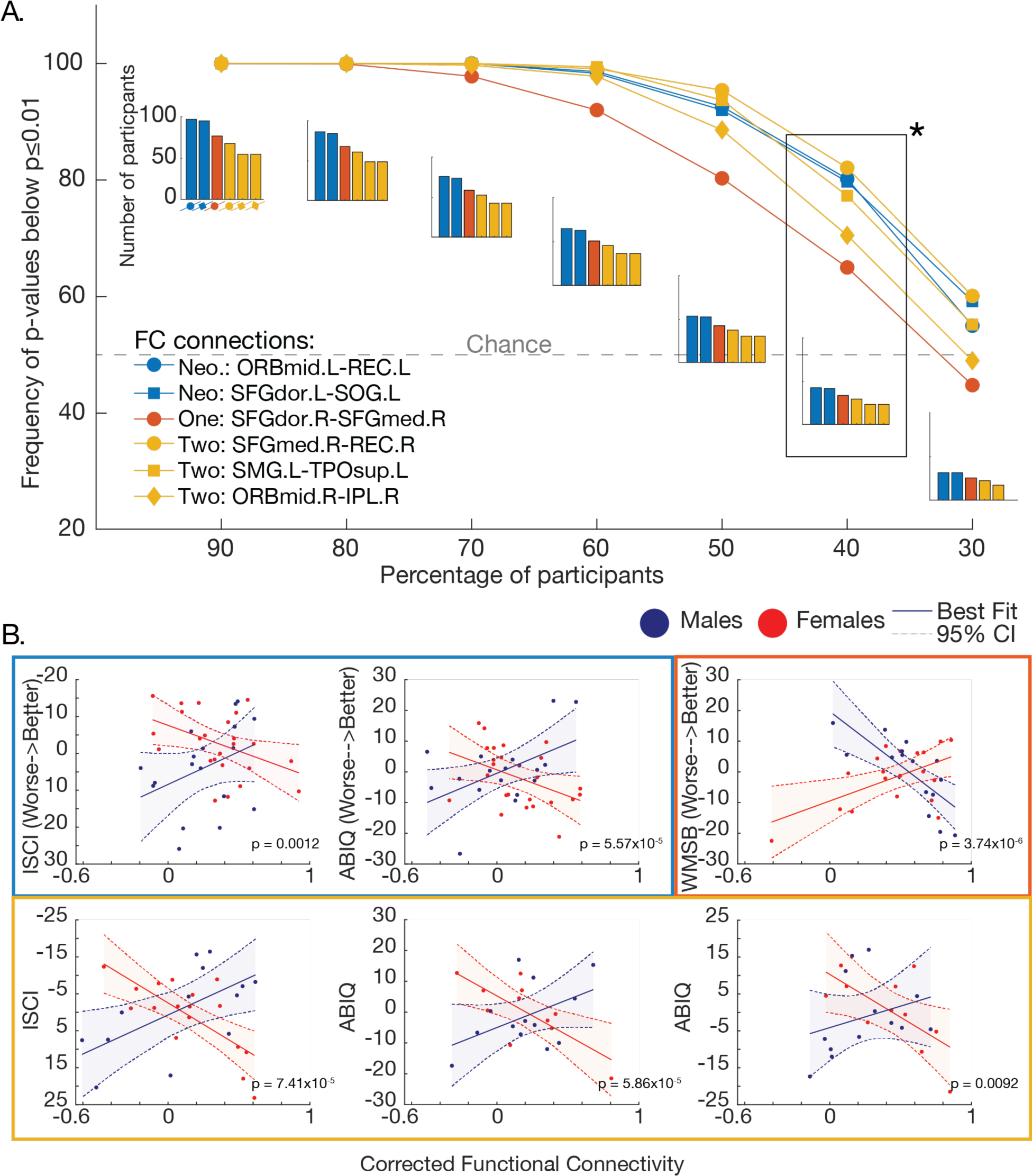
The consistency of functional connections across resampled subsamples. A. Datasets are resampled with replacement 1000 times across each subsample, a percentage (90-30%) of the original dataset size. The y-axis is defined by the probability of subsampled iterations that are significant (*p*≤.01). The grey line represents chance at 50% frequency of p-values below *p*≤.01. Bar graphs show the number of participants in each subsample. The asterisk (*) is presented in B. B. An example from one iteration of subsampling at 40% of the original dataset size. All interactions remain highly significant.

## 4. Discussion

This study revealed sex differences in brain-behavior relationships in the first two years of life by testing the interaction between sex and functional connectivity on future measures of behavior. In a sample of 316 participants at neonate, 1, and 2 years of age, we found six significant age-specific, intra-hemispheric connections with robust sex differences in functional connectivity associated with behaviors of inhibitory self-control, intelligence, and working memory. The connections occur within and between prefrontal regions across timepoints. Furthermore, these brain-behavior sex differences cannot be parsed from resting-state brain activity alone. These differences are only captured when behavior is also considered. The findings suggest that sex differences in brain-behavior relationships should be included as a factor in brain development heterogeneity.

Sex differences in early childhood cognition and behavior exist in abundance. However, evidence for the neural underpinnings for these processes is limited and not systematic. One of the main reasons for this discrepancy is that sex differences tend to be age-specific and vary throughout development. For example, in a review on sex differences in executive functioning, Grissom and Reyes describe the effect of age/timing as one of the many factors influencing the severity and expression of such sex differences [6]. Other studies have shown sex differences in cognitive [2–4] and socioemotional abilities [63,64] specific to the window of early childhood. For example, significant sex differences in communication (i.e., verbal and language skills) exist in preschool years [5,65–67] and can be predictive of later cognitive and clinical outcomes [65,68,69]. An area of missed opportunity is the potential sex differences in brain-behavioral relationships in early childhood. We previously used longitudinal data to show linear relationships between infant functional connectivity within temporal brain areas and childhood measures of language and intelligence [31]. In this current study, we expanded our investigation of brain-behavioral relationships to sex differences in whole-brain functional connectivity associated with measures of executive functioning (incl. working memory), intelligence, and anxiety. Remarkably, these unique connections were located within and between the prefrontal cortex (PFC).

The human PFC, which encompasses a significant portion of the frontal lobe, is considered a functionally unique, highly evolved brain region [70–72]. Pre- and postnatal factors, including sex differentiation, may influence the circuitry and development of the PFC [73]. Few studies have examined sex differences in fMRI connectivity before birth, but evidence suggests sex differences in prefrontal and frontal regions correlated to prenatal gestational age [74,75]. Mirroring our results in 2-year-olds, Wheelock and colleagues have reported fetal sex differences in the left temporal pole, a region with reciprocal connections to the orbitofrontal cortex within PFC [75]. After birth, prefrontal regions are the last to develop, with protracted development of the frontoparietal networks [29,61,76,77]. Synaptic pruning, the brain’s process of eliminating weak synapses [78], is slowest in the PFC [6] especially in the first years of life. This protracted development makes the brain more vulnerable to environmental perturbations [73,79,80]. Previously, we demonstrated a faster rate of growth from birth to 2 years of age in the frontoparietal network in boys [29]. In this study, we found sex differences in connectivity between the PFC and parietal regions in 2-year-olds (i.e., right inferior parietal lobule and supramarginal gyrus). These results imply differences that begin to emerge prenatally within the PFC, are retained through infancy.

The PFC has been implicated in higher cognitive behaviors such as those used in our study (i.e., executive functioning, intelligence, and anxiety). Sex differences have been widely reported in behavior. For example, weaker inhibitory control is often reported in boys in early childhood [2,81]. Females tend to score higher on intelligence tasks [3,4,82]; whereas a greater prevalence of males score at the lower end of these behavioral assessments. Subdomains of working memory also have been strongly correlated with intelligence in children and adults [83–85]. However, it is still unclear how these sex differences in behaviors relate to brain function. These reported sex differences in behavior may underlie the sex differences in PFC’s circuitry. For example, in adolescence through adulthood, the relationship between the PFC and task-based fMRI studies such as risk-taking [86,87], attentional switching [88,89], and interference inhibition [89], have shown differing functional activations in males compared to females. Using an independent component analysis on resting-state fMRI, Langeslag and colleagues demonstrated that functional connectivity in frontal and parietal regions associated with intelligence was significantly moderated by gender in children 6-8 years of age [90]. Moreover, this relationship between frontal-parietal regions and intelligence was stronger in girls. Collectively, our findings indicate that the functional connectivity mechanisms in infancy, particularly in the frontal lobes, may start to influence future behaviors shown to have sex differences well established in the literature.

Studies examining brain-behavior relationships in infancy are limited, but research in older children indicates a similar relationship to our results. In neonates and 2-year-olds, we showed that age-specific, intra-hemispheric connections displayed greater connectivity in males and lower connectivity in females in association with better behavioral scores. These intra-hemispheric locations might reflect the sex differences seen in a seminal study by Ingalhalikar et al., which demonstrated male structural connectivity as more intra-hemispheric and female connectivity as more interhemispheric in older children and young adults [91]. In conjunction, we showed that lower connectivity in males and greater connectivity in females were associated with worse behavioral scores in our current study. Rosch and colleagues observed stronger negative dorsolateral PFC fMRI connectivity associated with worse behaviors in ADHD, a condition impacting inhibition, in boys ages 8-12 [92]. Similar to our findings, Schmithorst and colleagues found that boys less than 9 years of age displayed positive correlations between regions associated with the frontal gyrus and intelligence [93]. However, no associations were found in girls under the age of 13. These age-related changes in cognitive function may be based on the peak of grey matter growth volume before adolescence [93–95]. Overall, the linear relationship found in boys and girls in our study resembles correlates between functional connectivity and behavior found in older populations.

In 1-year-olds, only one functional connection showed significant sex differences in brain-behavior relationships. This connection was located in the PFC between the right superior dorsal and medial gyri and displayed an opposite relationship to behavior compared to the connections at other age timepoints (Fig. 3B). We found lower connectivity in males and greater connectivity in females in association with better working memory. This relationship points to potential nonlinear developmental changes across age, reflective of our recent studies in functional connectivity in the first two years of life [30,31,96]. We previously investigated developmental fMRI trajectories in neonates through 2-year-olds and observed inter-subject variability as u-shaped across timepoints [96]. An overall decrease in variability in functional connectivity in the first year of life was found, followed by a region and age-specific increase in variability in the second year. Frontoparietal networks showed higher variability as well. This higher inter-subject variability across timepoints in functional connectivity may be indicative of greater sex differences between participants. Our findings highlight the age-specific connectivity of our cross-sectional analyses and the nonlinear relationship between age and functional connectivity over the first few years of life.

Evidence suggests that the genetic influence on sex differences can also affect the functional development of the PFC. Modification of male and female genetic expression begins before birth [16,97]. Fetal gonadal steroid hormones propel brain masculinization within a critical window in fetal development, with a maximal sex difference in testosterone concentration between 8-24 weeks gestation in humans [27,98]. The effect of these gonadal hormones leads to organizational effects on the developing brain with cascading adult consequences. For example, the SRY gene on the Y chromosome is a driver of testes development. Prenatal genetic variation of the SRY gene is potentially linked to the susceptibility to dopamine disorders in males [99–101]. Healthy dopamine levels enable successful cognitive control of the PFC [102–104]. Genetic studies have repeatably found alterations not only in the SRY gene but also in other genes involved with dopamine levels in ADHD [102,105,106]. Regardless of brain dysfunction, sex changes in gene expression identified in the PFC have been indicative of molecular differentiation in the male and female brains, particularly during infancy [107]. This strong relationship between genetics, sex differentiation, and the PFC in the prenatal and neonatal brain, implies a genetic basis for our data-driven results and its importance in brain-behavior relationships. Future work should consider gene-environmental interactions [108].

Sex differences in the functional connections found in this study were specific to brain-behavior relationships and not in functional connectivity alone. Our results may indicate that there are parallel mechanisms for sex differences with regards to brain functionality. One mechanism may demonstrate specific sex differences in brain functional connectivity alone, as shown in our previous work [31]. An alternative mechanism may be driven by contrasting brain-behavioral relationships, as shown here and in our other work presenting heterogenous brain-behavioral relationships in infancy [36]. Even though contrasting brain-behavioral relationships exist in these studies, when examining functional connections alone, connectivity patterns did not display any significant differences. This may reflect the ambiguity found in recent studies which have reported sex differences in the brain as sparse when taking brain and body volume into account [25,26,33]. One explanation for this inconsistency is that sex differences in the brain can be lost at group-level statistics when examining functional connectivity independently and disregarding subgroups that may exist in the population [41,109]. Taken together, heterogeneity in connectivity in the developing brain due to sex must be considered.

Our sex difference findings are robust against the reduction in sample size. Considering the previous report by Merk et al. claiming that brain wide associations with behavior stabilize at *N*≥2000 samples [109], we tested the robustness of our findings with step-wise reduction in the number of participants for each brain-behavior relationship detected. Surprisingly, even with a 60% reduction (i.e., 40% of the original sample), the observed significant sex differences in brain-behavioral relationships hold at *p*≤.01 level. These results stand in contrast to the previous report by Merk et al. and may suggest the advantage of well-defined, more homogeneous, single-site data [110]. Furthermore, our behavioral measures of have strong test-retest reliability and interrater reliability [36,38,47,55], which may have reduced inter-subject variability and increased consistency between brain-behavior relationships. The factors contributing to inconsistencies in multi-site detection of brain-behavioral relationships, may relate to potential heterogeneity in behavioral assessment instruments, scoring of assessments, and imaging data collection (both hardware and software).

While this study provides insight into sex as a factor in brain-behavioral relationships in infancy, there are limitations worth discussing. Environmental factors that also contribute to variation in our dataset are beyond the scope of this manuscript. Methodological factors such as scanning infants during natural sleep may be a potential confound [111]. However, monitoring of sleep stage using electrophysiological methods is challenging in infant populations [112]. Maternal education is another factor known to influence infant brain development and is an important determinant in childhood early development [113]. This factor was included as a covariate in our detection of brain-behavior associations but did not affect our results significantly. Another limitation is our use of parent-report executive function measures [55], which may also inherently contain a gender bias [114]. Finally, epigenetic changes are also due to prenatal and postnatal environmental exposures. However, these influences have been difficult to determine in group-level analyses [26].

## Perspectives and significance

Early childhood sex differences in cognition and behavior exist, but the neural underpinnings that give rise to such differences is relatively unknown. Here, we capture robust brain-behavior relationships distinct to sex in development. The developing brain undergoes the fastest rate of growth in volume across the human lifespan. Small brain differences can contribute to major changes in cognitive ability throughout childhood and adulthood [28,36,94]. During this time, abnormalities in brain structure and function increase the likelihood of developing future neurodevelopmental disorders and vulnerabilities. The age-specific functional connections found in this study provide insight into the PFC as a potential marker for later sex differences in cognition as well as neuropsychiatric disorders. Furthermore, such heterogeneity may be used to improve brain-based predictions. Subsequent studies should explore the replicability of these results as well as investigate the impact of genetics and environmental factors on the brain connections found in this study.

## 5. Conclusion

Our results underscore the importance of sex differences in brain-behavior relationships in the first two years of life. We found six age-specific, intra-hemispheric functional connections within the prefrontal cortex or the temporal pole. These functional connections displayed distinct brain-behavior mechanisms in males and females and emerge only when associated with future behavioral scores. Additionally, these brain-behavior relationships were robust against smaller sample sizes. Here, we highlight the importance of incorporating such heterogeneous relationships when examining functional connectivity in the developing brain. Even so, additional research is needed to explore the complex link between sex differences and brain-behavioral relationships due to the PFC’s involvement in genetics and environmental changes during infancy.

## 6. Declarations

### 6.1 Ethics approval and consent to participate

Parents or legal guardians of infants provided informed consent under protocols sanctioned by the University of North Carolina at Chapel Hill and Cedars-Sinai Institutional Review Boards.

### 6.2 Consent for publication

Not applicable.

### 6.3 Availability of data and materials

Processed data and code presented used for this manuscript is available on GitHub. Typically developing infants were part of the University of North Carolina (UNC) Early Brain Development Study, characterizing early childhood brain and behavior development. To protect study participants’ privacy, raw BOLD fMRI, meta, and behavioral data will not be shared. Processed data and code presented used for this manuscript is available on GitHub: https://github.com/sjfen/Sex_differences_brainbehavior_relationships_in_infancy.git

### 6.6 Authors’ contributions

**CRediT authorship contribution statement**

**Sonja Fenske:** Conceptualization, Investigation, Formal analysis, Writing – original draft, Visualization, Methodology. **Janelle Liu:** Writing – Review & Editing. **Haitao Chen:** Software, Data Curation. **Marcio A. Diniz:** Methodology **Rebecca Stephens:** Writing-Review & Editing, **Emil Cornea:** Data Curation. **John Gilmore:** Writing-Review & Editing, Resources, **Wei Gao:** Conceptualization, Methodology, Resources, Writing – review & editing, Supervision, Funding acquisition.

## 6.7 Acknowledgements

The authors thank the families who generously gave their time to participate in this study. We are grateful to the research assistants who collected and scored the 1-year-old, 2-year-old, and 4-year-old cognitive data and/or scored the Behavior Assessment System for Children–second edition and/or the Behavior Rating Inventory of Executive Function–Preschool version over the years for this study: Haley Parrish Black, Sadie Hasbrouck, Monica Ferenz Guy, Kassidy Jezierski, Margaret Hamilton Fox, Molly McGinnis, Mallory Turner, Emma Brink, Emily Bostwick, Margo Williams, Neha Patel, Portia Henderson, Jenna Obitko, Joe Blocher, and Rachel Steiner. The authors report no biomedical financial interests or potential conflicts of interest.

