## Supplemental Figure 1 for "Sex differences in brain-behavior relationships in the first two years of life"

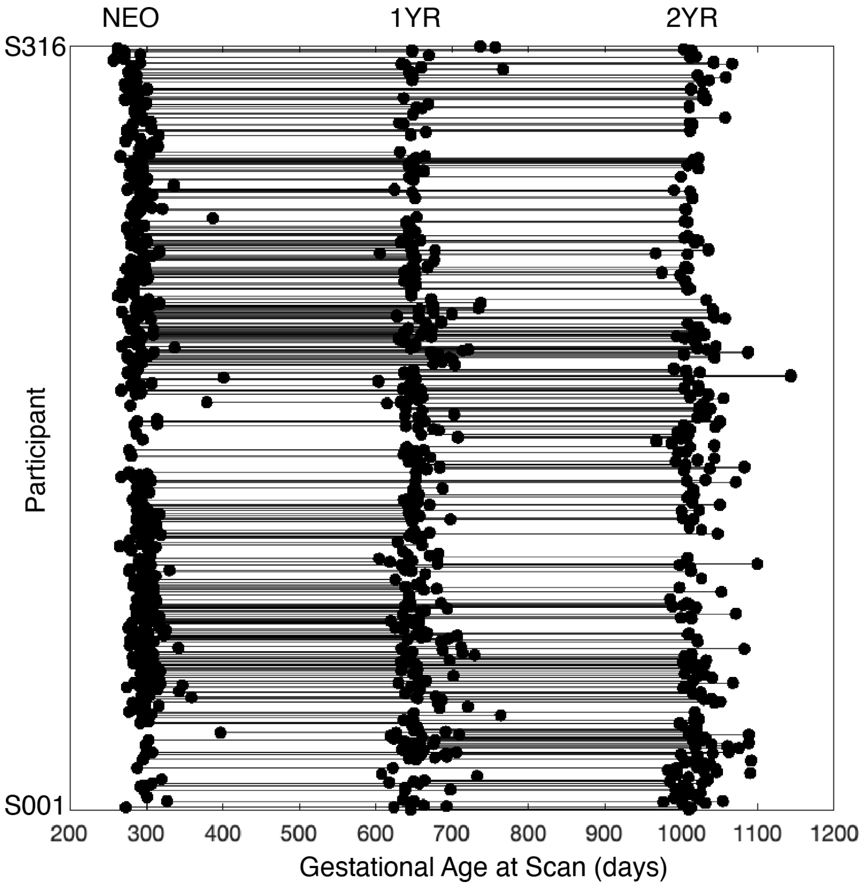


Figure S1. Data distribution. The distribution for all infant datasets with images passing quality control across gestational age at scan (*N*=316, with a total of 484 datasets). rsfMRI scans are represented at each dot and their longitudinal trajectories are represented by each line. Neonates (NEO): *n*=229; 1-year-olds (1YR): *n*=146; 2-year-olds (2YR): *n*=109; Neo and 1YR: *n*=85; 1YR and 2YR: *n*=60; NEO and 2YR: *n*=62; NEO, 1YR, and 2YR: *n*=38.
